# Mitochondrial bioenergetic signatures differentiate asymptomatic from symptomatic Alzheimer’s disease

**DOI:** 10.1101/2025.11.04.686626

**Authors:** Purba Mandal, Eugenia Trushina, Matthias Arnold, Rima Kaddurah-Daouk, Priyanka Baloni

**Affiliations:** School of Health Science, Purdue University, West Lafayette, IN 47907, USA; Purdue Institute for Integrative Neuroscience, Purdue University, West Lafayette, IN 47907, USA; PULSe Integrative Neuroscience program, Purdue University, West Lafayette, IN 47907, USA; Department of Neurology, Mayo Clinic, 200 First St. SW, Rochester, MN 55905, USA; Department of Molecular Pharmacology and Experimental Therapeutics, Mayo Clinic, 200 First St. SW, Rochester, MN 55905, USA; Institute of Computational Biology, Helmholtz Zentrum München – German Research Center for Environmental Health, Neuherberg, Germany; Department of Psychiatry and Behavioral Sciences, Duke University, Durham, NC, USA; Duke Institute of Brain Sciences, Duke University, Durham, NC, USA; Department of Medicine, Duke University, Durham, NC, USA

## Abstract

Asymptomatic Alzheimer’s disease (AsymAD) refers to individuals who, despite exhibiting amyloid-β plaques and tau pathology comparable to Alzheimer’s disease (AD), maintain cognitive performance similar to cognitively normal individuals. The resilience mechanism in these AsymAD individual remains understudied. We performed a systematic analysis comparing AsymAD and AD across multiple cohorts (ROSMAP, Banner and Mount Sinai), brain regions (BA6, BA9, BA36 and BA37) and neuronal and glial cell types using proteomics and transcriptomics data. AsymAD brains exhibited preserved mitochondrial bioenergetics, characterized by enhanced oxidative phosphorylation (OXPHOS), electron transport chain (ETC) activity, fatty acid and lipid metabolism, and branched-chain amino acid (BCAA) utilization. Pathways regulating mitochondrial complex biogenesis and calcium homeostasis were also upregulated. Key mitochondrial proteins such as MRPL47, CPT2, BCAT2, and IDH2, were consistently upregulated in AsymAD, whereas MACROD1 was downregulated. At the cellular level, excitatory neurons, including superficial, mid-layer, and deep-layer subtypes, exhibited the most preserved mitochondrial function, whereas vulnerable inhibitory subtypes, including PVALB and SST neurons, showed increased cellular abundance and bioenergetic activity. In contrast, microglia and oligodendrocytes proportions were reduced in AsymAD relative to AD. Our findings identify preserved mitochondrial bioenergetics as a defining feature of resilience in AD and suggest that enhancing NADH metabolism via NAD+ precursor-based interventions may potentially help in maintaining cognitive function despite amyloid and tau pathology.

## Introduction

Alzheimer’s disease (AD) is characterized by the accumulation of amyloid-β plaques and tau tangles^1,2^, which historically drove therapeutic strategies aimed at reducing these hallmark pathologies^3–7^. Despite decades of effort, most amyloid-targeted clinical trials, including Solanezumab (Eli Lilly)8, Aducanumab (Biogen)9–11, have failed, while for Lecanemab^10,12^ (Eisai/Biogen), the number of side effects is worse than the benefits offered, highlighting the need to explore alternative mechanisms that contribute to AD pathogenesis. Analysis of postmortem brain samples, positron emission tomography (PET) amyloid imaging studies and neuropsychological and memory binding assessments have identified a distinct group of individuals termed as the asymptomatic AD (AsymAD) cohort who exhibit plaque and tangle burdens comparable to AD patients yet maintain cognitive performance similar to controls/resistant cohort^13–15^. Unlike resistant individuals who resist pathology, these resilient individuals tolerate amyloid and tau accumulation without developing dementia. The mechanisms underlying this resilience remain poorly understood. Large-scale proteomic analysis using weighted gene co-expression network analysis (WGCNA) for brain and cerebrospinal fluid (CSF) samples has identified mitochondria as a key molecular module disrupted in AD compared to AsymAD, while transcriptomic analyses from separate studies have highlighted downregulation of specific mitochondrial genes such as *MT-CO1*, *MT-CO3*, *MT1G*, and *MT2A in* AD compared to resilient AD. However, the mechanisms by which mitochondria contribute to resilience in AsymAD individuals remain largely unexplored^15–17^.

Mitochondria are central to brain function, providing the energy required to sustain neuronal activity^18,19^. Each neuron can harbor hundreds to millions of mitochondria, reflecting the high metabolic demands of the brain^20,21^. Mitochondrial dysfunction impairs energy production, leading to neuronal death or compensatory degradation of brain tissue that could be contributing to the cortical atrophy observed in AD^22–24^. In contrast, asymptomatic individuals show preserved brain volume despite equivalent pathology^25,26^. Importantly, recent evidence suggests that mitochondrial dysfunction is not merely a consequence of AD but may act as a causal driver of neurodegeneration. Activation of mitochondrial G proteins has been shown to compensate for bioenergetic failure, leading to improved memory and reduced neurodegeneration in AD mouse model^27^. In humans, glucose hypometabolism precedes amyloid and tau deposition and may serve as a more sensitive diagnostic marker than classical pathology^28–31^. Other findings such as improved acylcarnitine profiles in resilient individuals^32^ and enhanced branched-chain amino acid (BCAA) utilization in APOE2 carriers relative to APOE4 carriers further support the role of bioenergetic resilience in mitigating AD risk^33^.

Although, several studies have independently identified mitochondria as a critical component in AD pathology, none have specifically focused on AsymAD individuals to investigate how mitochondrial features are preserved compared to AD across cohorts, brain regions, and cell types. To address this gap, we systematically analyzed proteomic and transcriptomic data to examine how mitochondrial regulatory and metabolic functions are maintained in AsymAD. We aimed to uncover mechanisms of mitochondrial resilience in AsymAD by identifying conserved mitochondrial features that may contribute to protection against neurodegeneration. Using proteomic data from the dorsolateral prefrontal cortex (BA9) across two cohorts (ROSMAP, and Banner), we identified mitochondrial pathways that are maintained in asymptomatic individuals. We also applied machine learning classifiers to extract a minimal set of mitochondrial proteins and trained logistic regression model to distinguish AD from AsymAD, validating their expression across four brain regions (BA6, BA9, BA36, BA37) and three cohorts (Banner, ROSMAP, Mount Sinai). To understand the cell type-specific differences between AD and AsymAD, we analyzed single-nucleus RNA sequencing and examined six major brain cell types: excitatory neurons, inhibitory neurons, astrocytes, microglia, oligodendrocyte precursor cells (OPC), and oligodendrocytes resolved into 14 subtypes. These results position mitochondrial bioenergetic preservation as a core feature of AsymAD pointing to new strategies for maintaining brain function despite amyloid and tau pathology.

## Methods

### Data acquisition

Proteomics and snRNA sequencing were obtained from the AD Knowledge portal from the Synapse database (https://www.synapse.org/). 4 brain regions were utilized from proteomics, dorsolateral prefrontal cortex (DLPFC; BA9) data was obtained from the consensus from 2 cohorts: ROSMAP (Religious Orders Study (ROS) involving older Catholic nuns, priests, and brothers across the United States; Memory and Aging Project (MAP) involving older people across Chicago metropolitan area) and Banner (Banner Sun Health Research Institute, Arizona), parahippocampal gyrus (PHG; BA36) was obtained from Mt Sinai (Mount Sinai School of Medicine Brain Bank, New York) and pre-motor cortex (BA6) and temporal cortex (BA37) was obtained from ROSMAP. For snRNA seq analysis, we leveraged postmortem brain data from ROSMAP cohort for all 6 cell types excitatory, inhibitory, astrocytes, microglia, OPC and oligodendrocytes. Synapse IDs are provided in the data availability section.

### Individual identification

We grouped the samples into control, AD, and AsymAD classes using the following rubric published by Johnson et al.^15^. Samples with CERAD 0–1 and Braak 0–3 without dementia at last evaluation were defined as control (if Braak equals 3, then CERAD must equal 0); cases with CERAD 1–3 and Braak 3–6 without dementia at last evaluation were defined as AsymAD; cases with CERAD 2–3 and Braak 3–6 with dementia at last evaluation were defined as AD. Individuals with dementia were defined based on their MMSE scores, specifically MMSE□<□24 or CDR□≥□1. The proteomics data consisted of a total of 488 samples in BA9 (106 control, 182 AD and 200 AsymAD), BA36 (45 control, 92 AD and 13 AsymAD), BA6 (25 control, 33 AD and 53 AsymAD), BA37 (25 control, 34 AD and 52 AsymAD). For snRNA seq analysis, we extracted a total of 347 samples consisting of 100 controls, 126 AD and 121 AsymAD cases.

### Mitocarta3 dataset

We extracted information from <u>MitoCarta3.0</u> dataset^34^, a comprehensive, curated resource that catalogs genes associated with mitochondrial structure, function, and biogenesis across various tissues and species. It includes over 1,100 mitochondria-associated genes identified through integrative approaches such as proteomics, mitochondrial targeting sequence predictions, and experimental validations. MitoCarta3.0 also provides tissue-specific expression profiles, enabling a deeper understanding of mitochondrial dynamics in different physiological and pathological contexts.

### Differentially expressed protein analysis

To identify the differentially expressed proteins in the dataset, we employed two approaches:

i. **Without cell type correction:** Differential expression was assessed across three comparisons, AsymAD vs AD, AD vs Control, and AsymAD vs Control, using two approaches. Proteomic data from the BA9 DLPFC was preprocessed, corrected for postmortem interval (PMI), age at death, and sex, with outliers removed, and values log-normalized using TAMPOR^15^. First, ANOVA followed by Tukey’s HSD test was applied to identify differentially expressed proteins. Second, the limma package in R(4.4.1) was used with Benjamini–Hochberg (BH) correction. In both approaches, proteins with an adjusted *p*-value < 0.05 were considered significant from our analysis. Mitochondrial proteins were identified using the MitoCarta 3.0 database.
ii. **With cell type correction:** To account for cellular composition, cell type marker proteins for neurons, astrocytes, microglia, oligodendrocytes, and endothelial cells were obtained from a published reference dataset ^15^. The expression matrix was aggregated by protein name to generate a protein × individual matrix of log-normalized expression values. For each sample, the mean expression of marker proteins for each cell type was calculated, yielding a matrix of cell type marker scores (rows = individuals, columns = cell types). These marker scores were then included as covariates in the limma design matrix prior to model fitting and BH correction. *design = model.matrix(∼ diagnosis + Astrocytes + Microglia + Neuron + Oligodendrocytes + Endothelia, data = cell_type_marker_scores)* Proteins with an adjusted *p*-value < 0.05 were considered significantly differentially expressed. Mitochondrial proteins were identified using the MitoCarta 3.0 database, consistent with prior analyses.

### Identifying features using machine learning models

Significant differentially expressed (DE) proteins identified from the proteomic analysis of the BA9 region were used as input for machine learning. Among the 258 DE proteins distinguishing AsymAD from AD, missing values from the expression data were imputed using k-nearest neighbor (KNN) imputation. Four classifiers: random forest^35^, gradient boosting^36^, recursive feature elimination (RFE)^37^, and LASSO-CV^38^ were applied to extract the most informative protein features. Proteins prioritized by these classifiers were then used to train a logistic regression model to classify individuals as AD or AsymAD based on protein expression. For RFE, the optimal number of features was determined by using the RFECV function and tested by plotting the mean cross-validated area under the curve (AUC) against the number of features. The best performance was achieved with 43 features, which were subsequently used to train the model. To avoid overfitting, 10-fold cross-validation was performed across all classifier models. Finally, SHAP (SHapley Additive exPlanations) analysis was conducted to interpret model predictions and identify the top 20 proteins most strongly contributing to discrimination between AD and AsymAD.

### Clubbing of cell types in single-nucleus RNA sequencing data

All eight cell-type objects were normalized and preprocessed by Mathys et al^39^. The original cell-type annotations from the published manuscript^39^ were retained and reformatted into more concise groupings in the table below following the conventions outlined in the subsequent publicationt^40^. Mic MKI67, T cells, CAMs were excluded from further analysis due to the limited number of cells precluding meaningful DEGs/pathway information. **Table 1** provides information on the cell clusters used for analysis.

**Table 1:**
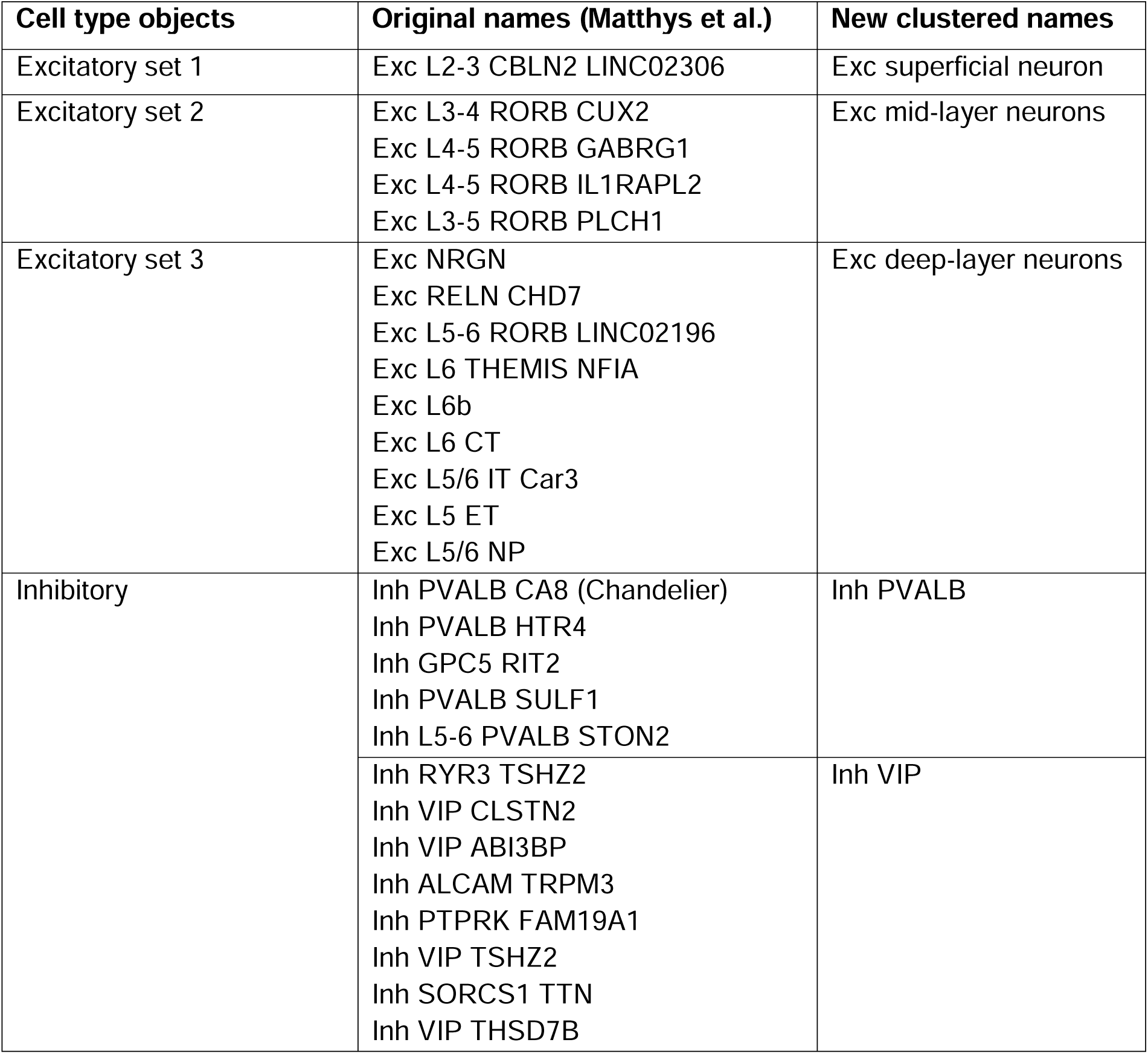
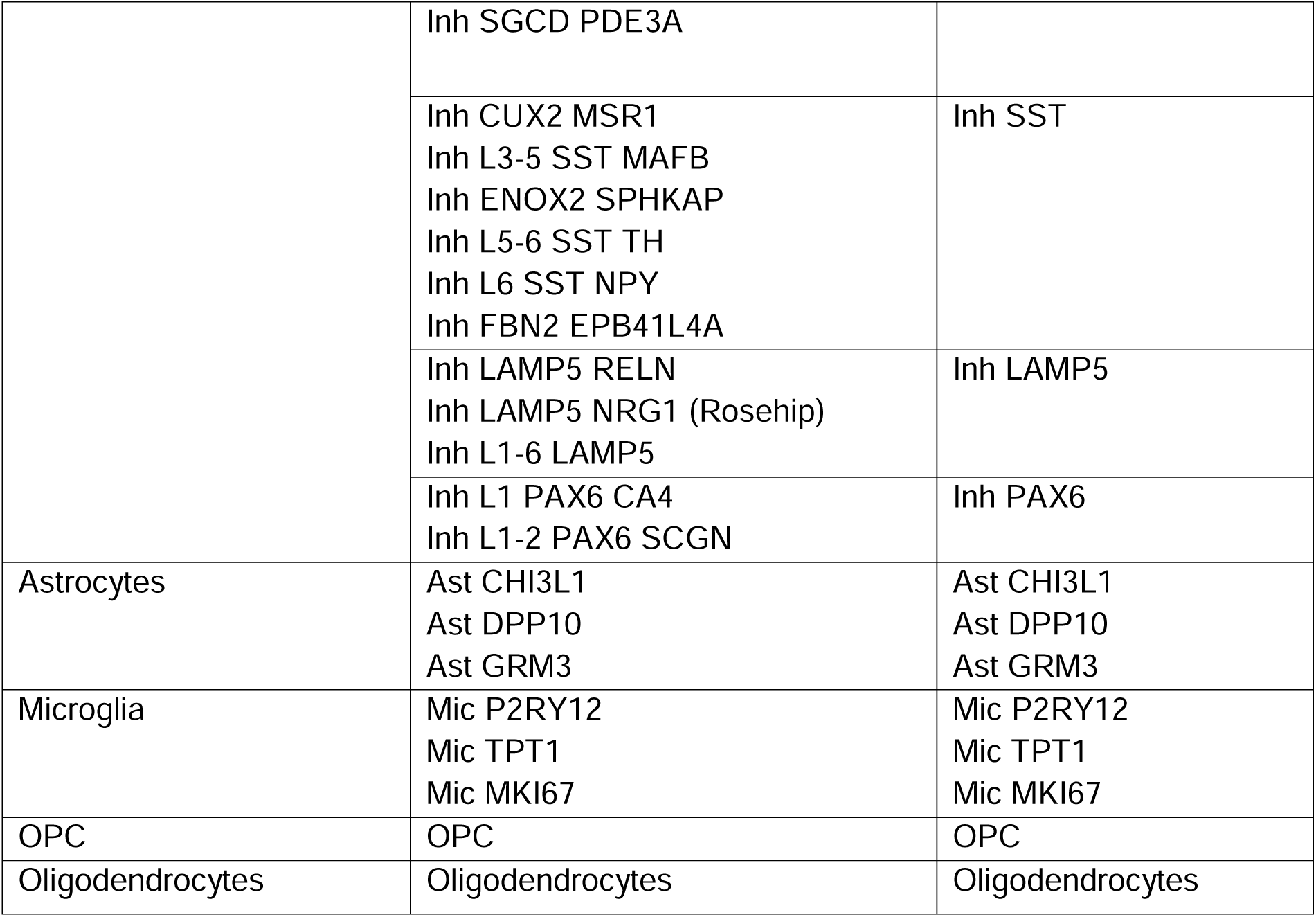
Cell cluster information used for study design and analysis.

### Differentially expressed gene analysis

Differential gene expression analysis was performed using the MAST package^41^ in R (4.4.1). A log fold-change threshold of 0.1 was applied, with the minimum expression cutoff set at *min.pct = 0.1* (i.e., genes expressed in at least 10% of cells in either group). Post-mortem interval (PMI), age at death, and sex were included as latent variables to account for potential confounding effects. Genes with a false discovery rate (FDR)– adjusted *p*-value < 0.01 were considered significantly differentially expressed. Mitochondrial genes were identified using the MitoCarta 3.0 database, consistent with previous analyses.

### Mitochondrial complexes mapping

Mitochondrial complexes I–V were mapped using the MitoCarta 3.0 database^34^. For the proteomics data, both nuclear- and mitochondrial DNA-encoded (MT) genes were retained for all complexes. For the snRNA-seq data, only nuclear-encoded mitochondrial complex genes were included, as nuclear genes were exclusively analyzed.

### Pathway enrichment analysis

We performed pathway enrichment analysis using the differentially expressed genes and proteins identified from our analysis as input and the Enrichr database (https://maayanlab.cloud/Enrichr/). We used KEGG Human 2021, Reactome Pathways 2024, and WikiPathways Human 2024 for all analyses. Significantly enriched pathways were identified based on adjusted p-values (< 0.05). The top pathways were ranked using the Combined Score, which integrates both statistical significance and the magnitude of enrichment, allowing comparison across multiple databases.

### Statistical analysis

Fisher’s exact test was applied to evaluate cell type abundance bias in the differentially expressed protein sets, both before and after cell type correction. Significant proteins were defined as those with an adjusted *p*-value < 0.05, while the background set included all proteins. For each cell type, a 2 × 2 contingency matrix was constructed consisting of: (i) marker proteins in the significant set, (ii) marker proteins not in the significant set, (iii) non-marker proteins in the significant set, and (iv) non-marker proteins not in the significant set. Fisher’s test *p*-values were generated for each cell type across all three comparisons (AsymAD vs AD, AD vs Control, AsymAD vs Control) and subsequently corrected for multiple hypothesis testing using the Benjamini–Hochberg (BH) procedure. Violin plots were generated for all brain regions (BA9, BA6, BA36 and BA37) to compare protein expression across Control, AD and AsymAD groups using *t*-test. The results were visualized using ggpubr() and ggplot2() functions in R (4.4.1).

## Results

### Study design for categorizing AD, AsymAD and Control samples

We included three groups in our study: Controls, Alzheimer’s disease (AD), and asymptomatic AD (AsymAD). We applied an integrated approach combining proteomics, machine learning, and single-nucleus RNA sequencing to identify mitochondrial genes/proteins that were differentially expressed between AD and AsymAD across cell types, brain regions, and cohorts (ROSMAP, Banner, and Mount Sinai) (**Fig. 1a**). In our classification, AsymAD cases were defined as individuals with CERAD scores 1–3 and Braak stages 3–6 but without dementia at their last evaluation, while AD cases had CERAD scores 2–3 and Braak stages 3–6 with dementia at last evaluation. Despite AsymAD cases exhibiting a similar pathological burden to AD, they had more brain weight (**Fig. 1b**) and maintained cognitive performance within the normal range, comparable to Controls as reflected by MMSE scores ≥24 (**Fig. 1c**).

**Figure 1:**
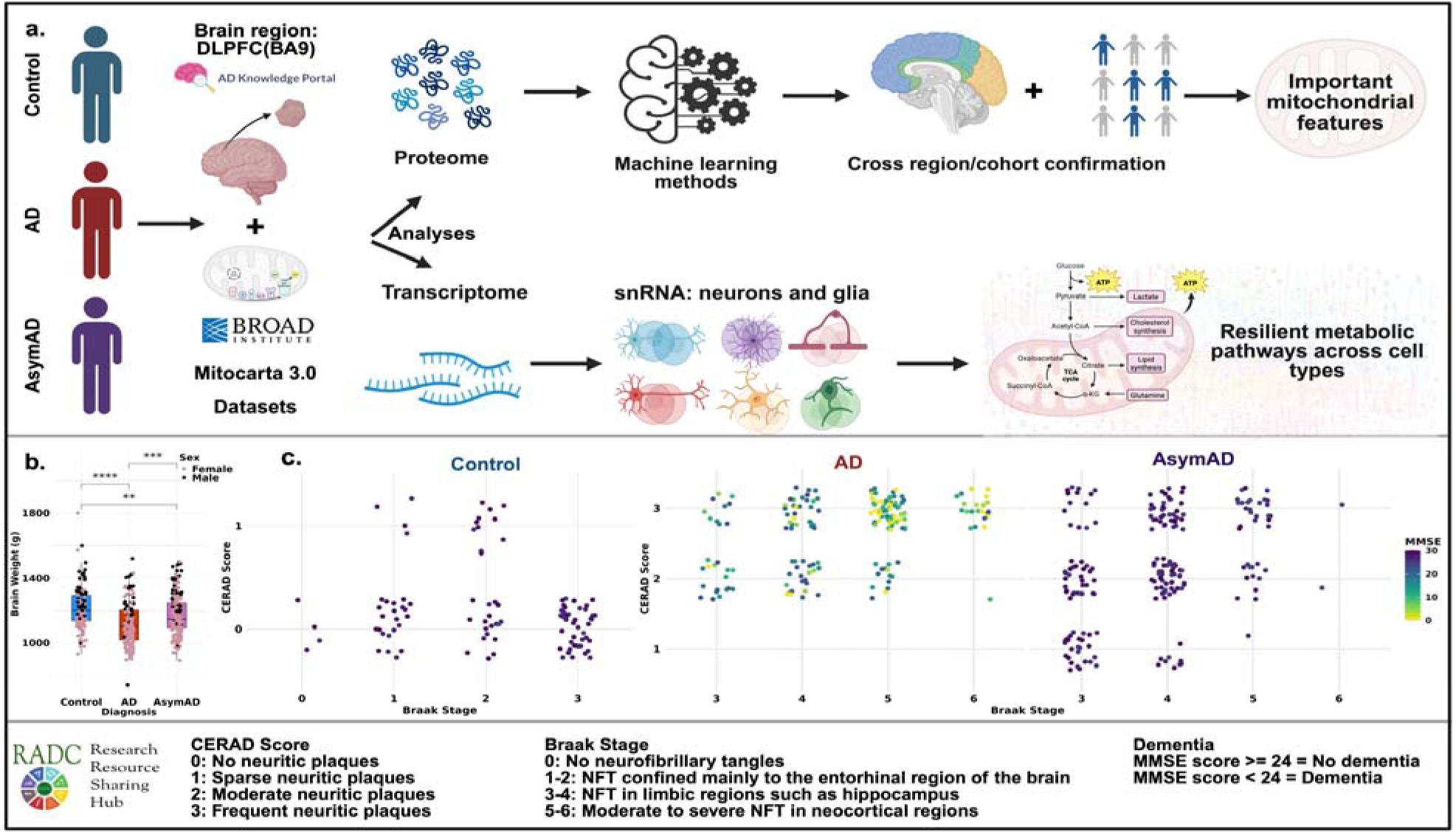
Asymptomatic AD (AsymAD) shows similar pathology to AD but similar cognition to Controls. a) Schematic for the overall theme of the paper, highlighting the 3 approaches used: proteomics, transcriptomics and machine learning to identify key mitochondrial proteins and pathways. b) Box plot showing the brain weight (in grams) for ROSMAP individuals (** = *p* < 0.01; *** = *p* < 0.001; **** = *p* < 0.0001). c) Scatter plot showing the Braak stages vs CERAD score for controls, AD and AsymAD colored by MMSE score per individual.

### AsymAD exhibit increased mitochondrial resilience compared to AD

To identify differentially expressed proteins (DEPs) between AsymAD vs AD, we first calculated the bias due cell type abundance, given that AsymAD brains retain cognition and show minimal brain atrophy despite the presence of neuropathological hallmarks (**Fig. 1b**). Using established marker proteins for neurons, astrocytes, microglia, oligodendrocytes, and endothelial cells reported by Johnson et al.^15^, we observed that AsymAD brains exhibited higher neuronal abundance, but lower levels of microglia, oligodendrocytes, and endothelial cells compared to AD (**Supplementary Fig. 1a,b**).

To control for these differences and capture true protein-level changes, we applied the limma workflow, incorporating cell type marker scores as covariates in the design matrix. We further validated the correction using Fisher’s exact test to assess cell type enrichment before and after adjustment. We observe that the initial enrichment of neuronal, microglial, and endothelial proteins in uncorrected data was removed post-correction, confirming that the final set of DEPs reflects protein levels independent of cell-type composition (**Supplementary Fig. 1ci-iii**).

To explore the biological processes associated with 1757 DEPs obtained after cell type correction, we performed pathway enrichment analysis using KEGG, Reactome, and WikiPathways databases. For the pathways associated with upregulated DEPs in AsymAD vs AD, we observed enrichment for metabolism, translation, the TCA cycle, and ETC, suggesting that AsymAD individuals retain metabolic, mitochondrial and translational activity better than AD (**Supplementary Fig. 1d**). In contrast, pathways associated with downregulated DEPs in AsymAD vs AD showed enrichment for AUF1-mediated RNA destabilization, translational inhibition, and components of the innate immune system, indicating a potential shift toward enhanced protein synthesis and reduced inflammatory response in AsymAD (**Supplementary Fig. 1e**). 258/1757 DEPs mapped to the MitoCarta 3.0 database, indicating significant alterations in mitochondrial protein expression in AD compared to AsymAD. To further validate the robustness of our cell type-corrected pipeline, we used ANOVA with Tukey’s HSD and limma without cell-type correction to identify mitochondrial DEPs across diagnoses. 221 DEPs were commonly identified across all three approaches in AsymAD vs AD. For AD vs Control, we identified 198 overlapping DEPs, confirming that our strategy yielded a reliable and conservative set of mitochondrial proteins (**Supplementary Fig. 1f–h**).

Out of the 258 overlapping DEPs between cell type corrected data and MitoCarta 3.0, 229 DEPs were upregulated and 29 downregulated in AsymAD vs AD (**Fig 2a**). Pathway enrichment analysis for these 229 upregulated proteins resulted in mitochondrial regulatory as well as metabolic pathways. The top enriched metabolic pathways included ETC, OXPHOS, the TCA cycle, BCAA, fatty acid and propanoate metabolism. For regulatory pathways, the top enriched pathways included biogenesis of Complexes I, III, and IV, mitochondrial translation, protein import and degradation, fusion and fission regulation and mitochondrial calcium ion transport (**Fig 2b and c**).

**Figure 2:**
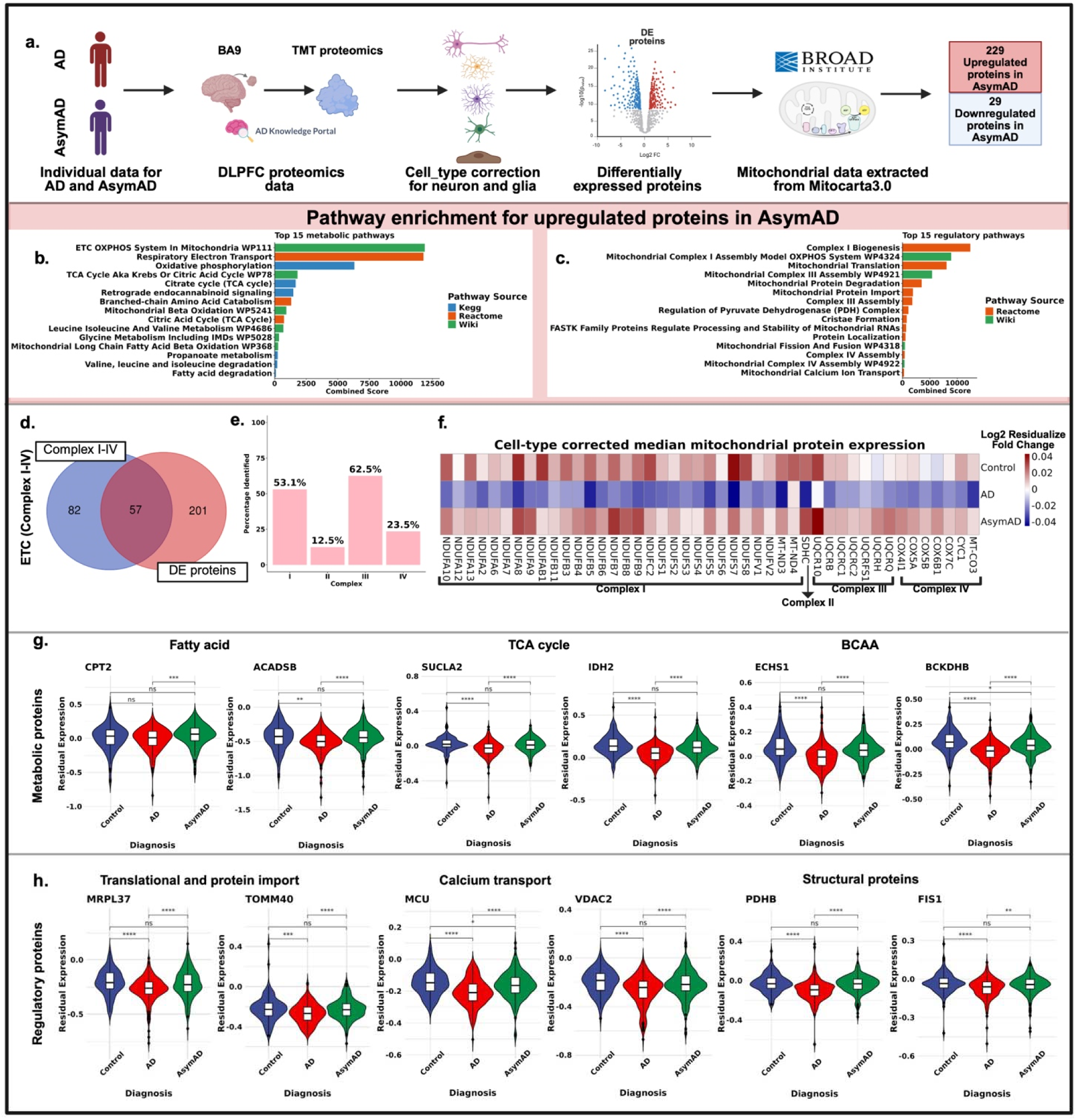
AsymAD maintain mitochondrial bioenergetics (ETC, the TCA cycle, BCAA and fatty acid) compared to AD. a) Schematic diagram representing 229 upregulated and 29 downregulated mitochondrial proteins between AsymAD vs AD using BA9 proteomics data after correcting for cell type bias. b & c) Pathway enrichment analysis shows upregulation of metabolic and regulatory mitochondrial pathways in AsymAD vs AD. d) Venn diagram showing the number of mitochondrial proteins overlapping with mitochondrial OXPHOS Complexes I-IV. e) Percentage bar plot of overlap for the DEPs across mitochondrial Complexes. f) Residualized median expression of protein heatmap of mitochondrial Complexes across Control, AD and AsymAD individuals after cell type, PMI, age of death and sex correction. g i-vi) Violin plot of key proteins involved in fatty acid, the TCA and BCAA across Control, AD and AsymAD using *t*-test. h i-vi) The Violin plot of key proteins involved in regulatory mitochondrial functions: Translation, calcium transport and structural proteins across Control, AD and AsymAD using *t*-test. (ns = *p* > 0.05; * = *p* < 0.05; ** = *p* < 0.01; *** = *p* < 0.001; **** = *p* < 0.0001)

Since OXPHOS was the most enriched pathway in AsymAD, we examined the overlap of our dataset with proteins reported in the ETC. In total, 57 proteins overlapped with ETC complexes (**Fig. 2d**). We found the following % overlap with the complexes: 53.1% for Complex I, 12.5% for Complex II, 62.5% for Complex III, and 23.5% for Complex IV. We observed the highest coverage accounting for over half of the proteins in Complexes I and III (**Fig. 2e**). We compared protein expression for mitochondrial complexes across AD, AsymAD, and Controls, and observed that AsymAD and Controls shared similar expression profiles across all four Complexes, whereas AD showed consistent downregulation (**Fig. 2f**). This finding supports our observation that OXPHOS is the most conserved pathway in AsymAD. The heatmap displays fold change values for proteins in Complexes I-IV after correcting for cell type abundance, highlighting true protein abundance changes rather than shifts driven by cellular composition.

Other than OXPHOS, we also examined key metabolic and regulatory proteins that are essential for energy generation. Proteins involved in fatty acid oxidation (CPT2; *Carnitine Palmitoyltransferase 2*, ACADSB; *Acyl-CoA Dehydrogenase, Short/Branched Chain*), BCAA metabolism (ECHS1; *Enoyl-CoA Hydratase, Short Chain 1*, BCKDHB; *Branched-Chain Ketoacid Dehydrogenase E1 Subunit Beta*) and the TCA cycle (SUCLA2; *Succinate-CoA Ligase ADP-Forming Beta Subunit* and IDH2; *Isocitrate Dehydrogenase 2*) were significantly upregulated in AsymAD vs AD (**Fig. 2g**). In addition, calcium regulatory proteins (MCU; *Mitochondrial Calcium Uniporter*, VDAC2; *Voltage-Dependent Anion Channel 2*), mitochondrial translation factors (MRPL37; *Mitochondrial Ribosomal Protein L37*, TOMM40; *Translocase of Outer Mitochondrial Membrane 40*), and structural components (PDHB; *Pyruvate Dehydrogenase E1 Beta Subunit*, FIS1; *Fission 1 Homolog*) were also upregulated in AsymAD, with expression levels resembling those of Controls (**Fig. 2h**). All up and down regulated mitochondrial proteins are provided in **Supplementary Table A**.

### Mitochondrial features associated with fatty acid, the TCA cycle, and amino acid metabolism distinguish AsymAD from AD

Using the 258 DEPs for AsymAD vs AD in the BA9 region, we developed a machine learning model to identify the most informative features that distinguish AsymAD from AD individuals. Protein expression data were imputed using KNN (k-nearest neighbors), and proteins with >15% missing values were plotted. KNN imputation did not introduce major shifts in mean or variance and preserved the overall distribution of protein expression (**Supplementary Fig. 2a, b**). Following imputation, we applied four classifiers, namely Random Forest, Gradient Boosting, RFE (Recursive feature elimination), and LASSO-CV, and features selected by each were used to train a logistic regression model with a 10-fold cross-validation (**Fig. 3a**). Among the classifiers, RFE selected 43 features (**Supplementary Fig. 2c, d**) and achieved the best performance (train AUC = 0.92, validation AUC = 0.88, accuracy = 0.81) (**Fig. 3c**). In addition, we also tested the combined intersection of features identified by all four classifiers (*n* = 16; **Fig. 3b**, black outline). A full list of features selected by each model is provided in **Supplementary Table B.** To assess biological relevance, we compared the 43 RFE-selected proteins against previously nominated AD-associated proteins from AGORA (ref). Six proteins overlapped (**Fig. 3b**, pink outline), of which four proteins, ALDH4A1, DBT, DAP3, and IDH2, were present in the top 20 SHAP features (**Fig. 3d**). SHAP analysis revealed that higher expression of MRPL47, FTH1, BDH1, and CPT2 was predictive of AsymAD, whereas greater expression of MACROD1, ALDH9A1, DAP3, and FASN was associated with AD. Notably, several proteins involved in BCAA and fatty acid metabolism were enriched among predictors of AsymAD (**Supplementary Fig. 2e**).

**Figure 3:**
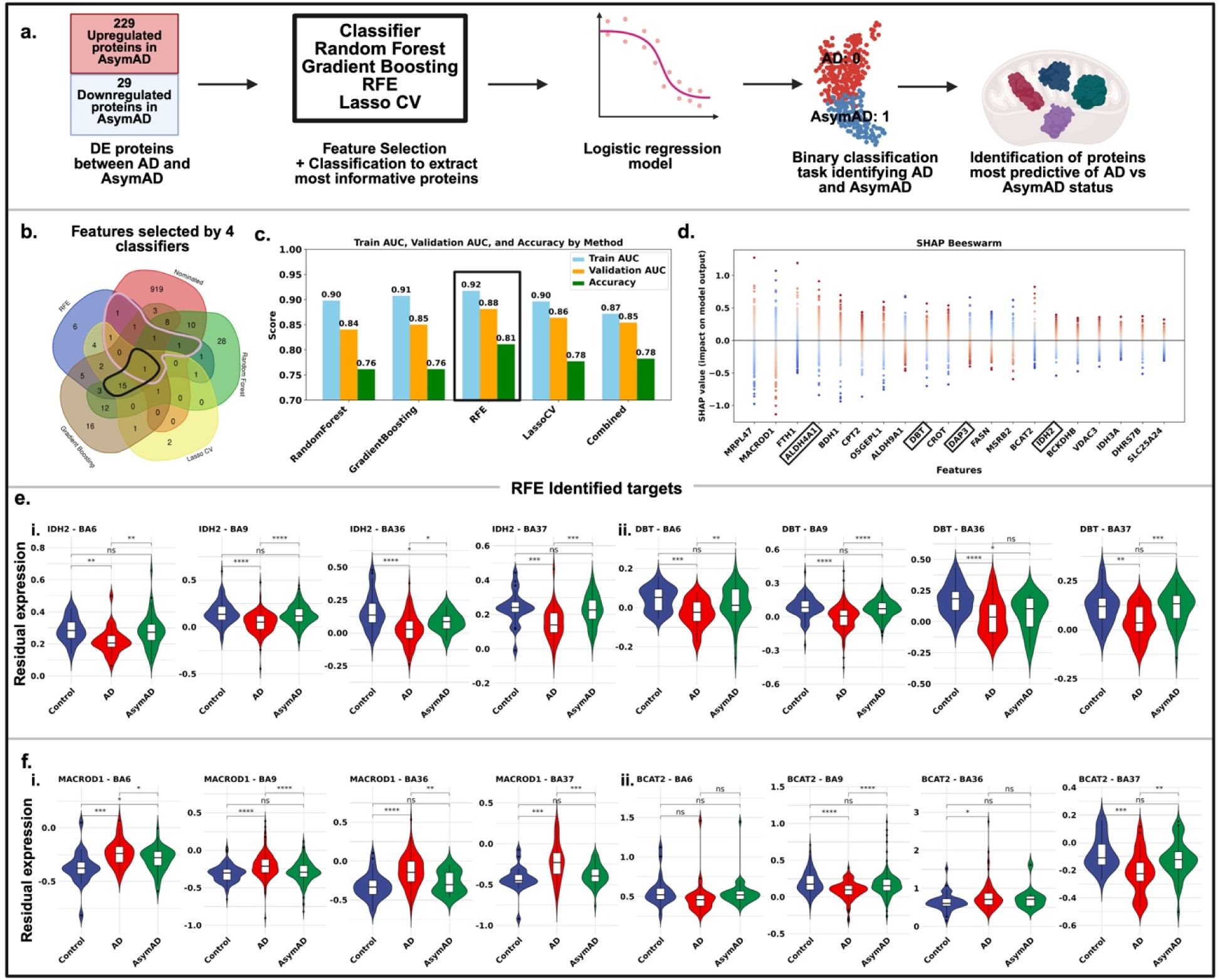
Identification of the top 20 mitochondrial features that distinguish AsymAD from AD. a) Schematic of different classifiers used to identify key mitochondrial features from the 229 upregulated and 29 downregulated proteins between AsymAD vs AD identified using BA9 proteomics data. b) The Venn diagram of common proteins identified by each classifier and nominated proteins (pink = common nominated proteins with RFE; black = common protein). c) Bar plot reporting the train AUC, validation AUC and accuracy for each classifier-model, including the common proteins from all classifiers. d) SHAP plot of the top 20 proteins used by the RFE-logistic model to distinguish AsymAD from AD. e) The Violin plot of nominated targets + identified by RFE, IDH2 and DBT proteins across 4 brain regions (statistical significance calculated using *t*-test). f) Violin plot of identified proteins by RFE, MACROD1 and BCAT2 across 4 brain regions (statistical significance calculated using *t*-test) (ns = *p* > 0.05; * = *p* < 0.05; ** = *p* < 0.01; *** = *p* < 0.001; **** = *p* < 0.0001)

Finally, to test the robustness of these features, we examined their expression across additional cohorts and brain regions. IDH2 (*Isocitrate Dehydrogenase 2*) and DBT (*Dihydrolipoamide Branched Chain Transacylase E2*) involved in the TCA cycle and BCAA metabolism was consistently upregulated in AsymAD compared to AD across BA6, BA9, BA36, and BA37 (**Fig. 3e**), whereas MACROD1 (*Mono-ADP-Ribosylhydrolase 1;* a post-translational modification involved in DNA repair, transcription, and stress response) was consistently downregulated in AsymAD across the same regions and BCAT2 (*Branched-Chain Aminotransferase 2*) upregulated in BA9 and BA37 (**Fig. 3f**). Among the 43 features selected by RFE, several solute carrier mitochondrial family (SLC25) proteins were upregulated in AsymAD, including SLC25A46 (mitochondrial dynamics and lipid transfer) and SLC25A27 (uncouples oxidative phosphorylation). We also observed upregulation of DAP3 (*Death-Associated Protein 3 (Mitochondrial Ribosomal Protein S29*)) and FIS1 (*Fission 1 Homolog*), essential for mitochondrial maintenance (**Supplementary Fig. 2f–g**).

### Mitochondrial bioenergetic resilience across brain cell types in AsymAD

snRNA-seq data from the BA9 brain region were analyzed, with individuals classified as Control, AD, or AsymAD, based on previously established criteria published in Johnson et al.^15^(**Supplementary Fig. 3a**). Across 1.9 million cells, six major cell types were considered: excitatory neurons, inhibitory neurons, astrocytes, microglia, oligodendrocyte precursor cells (OPCs), and oligodendrocytes. The original snRNA-seq dataset contained a large number of subtypes, including 14 excitatory neuronal subtypes, 25 inhibitory neuronal subtypes, 3 astrocytic subtypes, and 5 immune populations. To reduce complexity while retaining biological meaning, these were consolidated into broader categories (refer to the methods section: Clubbing of cell types in single-nucleus RNA sequencing data). Specifically, excitatory neurons were grouped into superficial, mid-layer, and deep-layer excitatory neurons, while inhibitory neurons were grouped into PVALB, VIP, SST, and LAMP5 subtypes (**Supplementary Fig. 3b**). Other populations, including astrocytes, microglia, OPCs, and oligodendrocytes, were kept as originally defined in Mathys et al^39^.

We observed comparable proportions of male and female samples, as well as diagnosis groups (Control, AsymAD, and AD individuals) (**Fig. 4a**). To assess whether shifts in cell type composition occurred across disease progression, we calculated the proportion of each cell type subclass within each group. As expected, the inhibitory neuronal subtypes Inh SST and Inh PVALB were reduced in AD, consistent with their known vulnerability, but were retained in AsymAD (**Fig. 4b**). We observed an interesting pattern for microglia and oligodendrocytes showing progressive increases in cell number from control to AsymAD to AD, consistent with previous reports of microglial expansion and oligodendrocyte enrichment during AD progression^39,42,43^. Similar result was observed for oligodendrocytes from the cell type resolved proteomics data analysis performed earlier (**Supplementary Fig 1b**). Remaining cell types showed trends that did not reach significance (**Supplementary Fig. 3c**).

**Figure 4:**
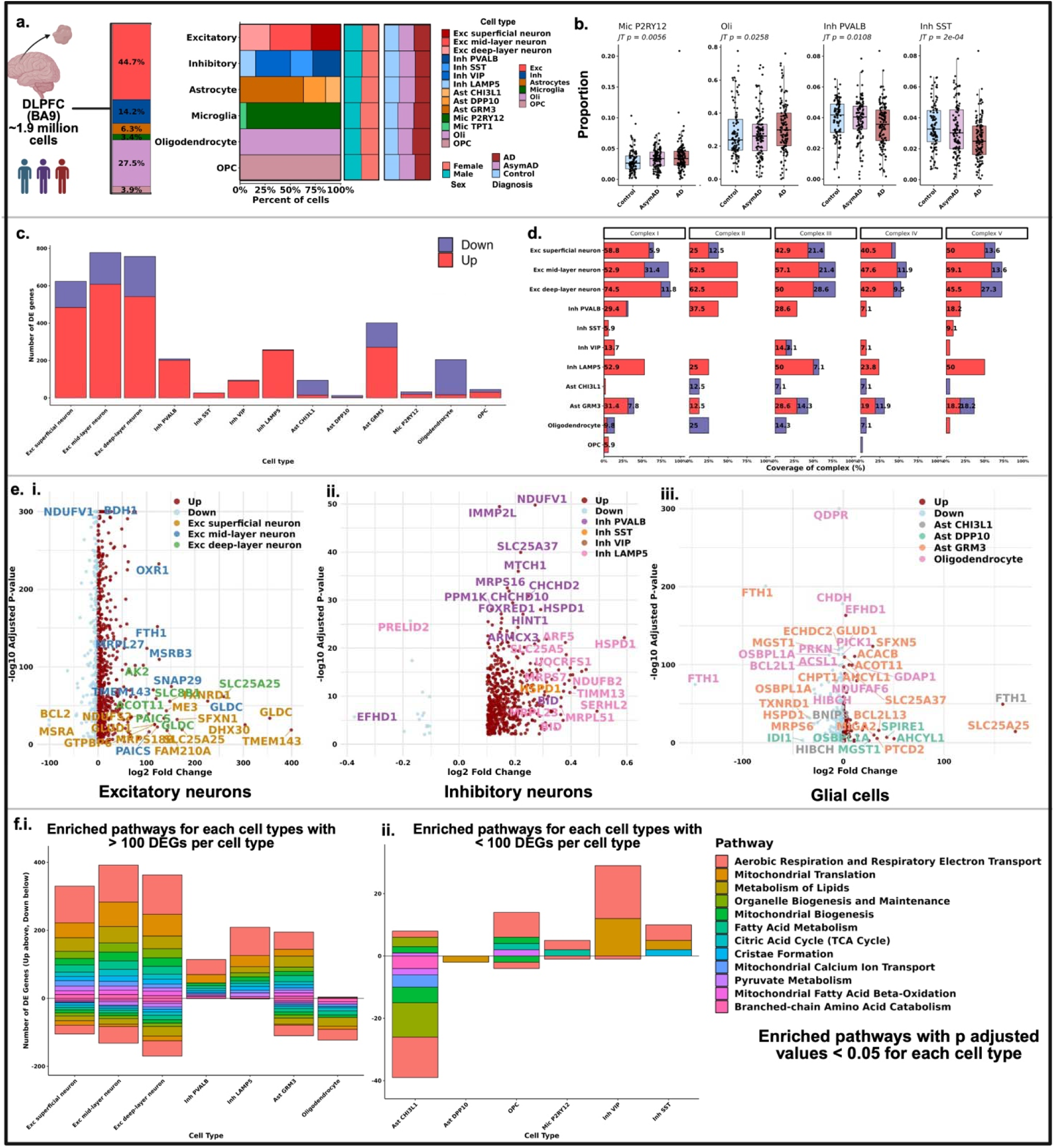
Cell type-specific mitochondrial changes in AsymAD vs AD. a) BA9 DLPFC from Control, AsymAD and AD used for analysis with equal proportions of male and female samples clubbed into 6 main types and 14 subtypes. b) Box plots of proportion of each cell type per diagnosis to identify a trend when going from Control AsymAD AD using Jonckheere’s trend test. c) Bar plot of up- and down-regulated DEGs between AsymAD vs AD obtained using MAST analysis per cell subtype. d) Bar plot reflecting the coverage of nuclear encoded genes of mitochondrial Complex present per cell subtype. e) Volcano plot showing the top DEGs across cells color coded by up and downregulation and subtype; i) DEGs for excitatory neuronal subtypes (superficial, mid-layer and deep-layer neurons); ii) DEGs for inhibitory neuronal subtypes (PVALB, VIP, SST and LAMP5); iii) DEGs for glial cell subtypes (Ast CHI3L1, Ast DPP10, Ast GRM3 and oligodendrocytes). fi) Bar plot showing the top enriched pathways associated with greater than 100 DEGs per cell subtype, ii) bar plot showing the top enriched pathways associated with less than 100 DEGs per cell subtype.

To investigate cell type-specific changes between AsymAD and AD, we performed MAST differential expression analysis (**Fig. 4c**). The largest number of mitochondrial DEGs was observed in excitatory neurons, the majority of which were upregulated in AsymAD compared to AD, followed by astrocytes and inhibitory neurons. In contrast, oligodendrocytes uniquely exhibited a higher proportion of downregulated genes, differing from the upregulation trends seen in other cell types (**Fig. 4c**). Given that OXPHOS was the most preserved pathway in AsymAD (**Fig. 2b**), we next assessed the coverage of mitochondrial Complexes I-V for DEGs (both up- and downregulated in AsymAD vs AD) across cell types (**Fig. 4d**). Excitatory neurons showed the highest coverage, especially the upregulated genes in mid-layer and deep-layer neuronal subtypes mapping to mitochondrial complexes. Among inhibitory neuron subtypes, predominantly upregulated genes in PVALB and LAMP5 neurons, followed by VIP and SST mapped to the mitochondrial Complexes. In astrocytes, the homeostatic population (Ast GRM3 ^44^) displayed the strongest coverage of Complexes I & III, whereas activated astrocytes (Ast CHI3L1 ^45^) exhibited higher coverage of Complex II, driven largely by downregulated genes, indicating reduced activity compared to homeostatic astrocytes. Oligodendrocytes showed more coverage of Complex II for genes that were mostly downregulated, while OPCs demonstrated upregulation of Complex I genes. These results indicate broad preservation of mitochondrial bioenergetics across cell types in AsymAD, with activated or inflammatory astrocytes and oligodendrocytes showing reduced activity, likely reflecting lower metabolic demands within their environment.

To investigate the most significantly differentially expressed genes (DEGs) across cell subtypes, we generated volcano plots for the three major cell classes: excitatory neurons, inhibitory neurons, and glial cells (**Fig. 4e.i–iii**). In excitatory neurons, *NDUFV1, BDH1, BCL2, TREM143, GLDC*, and *SLC25A25* emerged as top DEGs between AsymAD vs AD. For inhibitory neurons, prominent DEGs included *NDUFV1, EFHD1, PRELID2, SLC25A37,* and *IMMP2L*. For glial cells, *QDPR, FTH1,* and *SLC25A25* were the most significantly altered genes. Several of these genes overlapped with the post-translational protein features identified by RFE, suggesting consistency between transcriptional and proteomic signatures of mitochondrial resilience. To determine which pathways were associated with the DEGs, we performed Reactome pathway enrichment analysis for each subclass of cells (**Fig. 4fi-ii).** Across most cell types, we observed upregulation of metabolic pathways associated with the upregulated DEGs, including ETC, lipid and fatty acid metabolism, and the TCA cycle, as well as regulatory pathways such as mitochondrial translation, calcium ion transport, and cristae formation. Oligodendrocytes were the only cells that predominantly exhibited downregulated pathways. Similarly, activated astrocytes marked as Ast CHI3L1 displayed marked downregulation of metabolic pathways compared to the homeostatic astrocytic population (Ast GRM3). A full list of all DEGs across all cell types are provided in **Supplementary Table C**.

To investigate why oligodendrocytes uniquely exhibited downregulation in AsymAD, we compared this cell type across all three conditions: AD vs Control, AsymAD vs AD, and AsymAD vs Control (**Supplementary Fig. 3d**). We observed a distinct pattern especially for mitochondrial metabolic pathways that were upregulated in AD vs Control and AsymAD vs Control, indicating higher mitochondrial activity in the presence of pathology compared to no pathology. However, in AsymAD vs AD, these pathways were downregulated, suggesting that oligodendrocyte mitochondrial activity is selectively lower in the AsymAD brain environment relative to AD.

### Identifying small-molecule compounds associated with bioenergetic balance in AsymAD

Using the 258 differentially expressed proteins identified in Fig. 1, we queried DrugBank to identify small-molecule compounds associated with these proteins. NetworkAnalyst analysis identified 45 druggable proteins corresponding to 16 candidate drugs or small molecules. To assess the biological and pathological relevance of these interactions, we integrated data from AD Atlas^46^ (https://adatlas.org/) compiling protein–metabolite associations, their relationships to AD-related phenotypes (including cognition, tau pathology, and global pathology burden), and small-molecule interactions into a unified network to identify hub nodes. Among these, BDH1, IDH3A, and ALDH9A1 proteins, identified from convergence across machine learning, snRNA-seq, and proteomics analysis (highlighted as bold and represented in the Venn diagram in (**Fig. 5**)), were found to interact with dihydroorotate and hydroxylysine. Many proteins in this network were linked to NAD⁺/NADH-dependent redox reactions, highlighting the centrality of mitochondrial redox balance in AsymAD bioenergetic resilience. While NADH itself doesn’t cross the blood-brain barrier, these findings emphasize that modulating brain NAD^+^ metabolism could restore redox equilibrium and support mitochondrial function. These results highlight a convergent network of proteins, metabolites, pathology, and druggable targets that collectively point to mitochondrial NAD^+^/NADH metabolism as an important feature of bioenergetic stability in AsymAD (**Fig. 5**).

**Figure 5:**
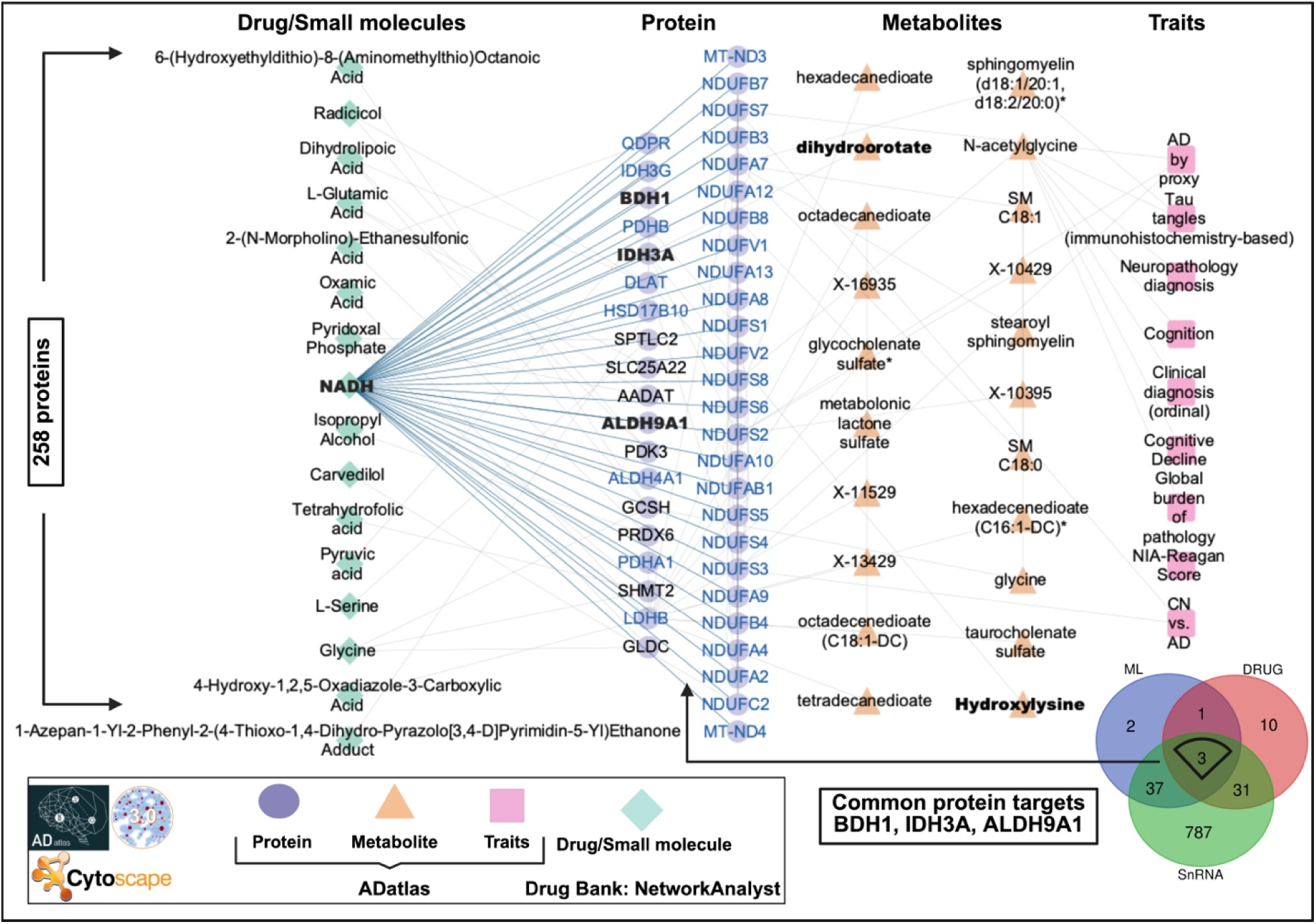
Integrated network of proteins, metabolites, AD pathology features, and linked drugs/small molecules.

We further characterized how each of the 16 small molecules interact with broader protein networks using DrugBank via BioGRID and annotated their pathways through Reactome. NADH exhibited the highest number of interacting protein partners (144) (**Supplementary Fig. 4a**), primarily enriched in pathways related to the electron transport chain (ETC), Complex I biogenesis, and mitochondrial protein degradation (**Supplementary Fig. 4b**). L-glutamic acid followed with 69 interacting partners (**Supplementary Fig. 4a**), involved in neurotransmission and synaptic signaling. Across all identified compounds, ‘metabolism of amino acids and derivatives’ emerged as the most significantly targeted pathway, highlighting the importance of metabolic regulation in the context of AsymAD (**Supplementary Fig. 4b**). Comprehensive pathway and protein-compound interaction details are provided in **Supplementary tables D and E**. This convergence of protein, metabolite, and small molecule interactions highlights redox modulation as a potential metabolic strategy to sustain neuronal function and counteract Alzheimer’s disease progression.

## Discussion

In this systematic study, we leveraged proteomics and transcriptomics from post-mortem brain samples and developed machine learning models to identify important metabolic features that distinguish asymptomatic (AsymAD) from symptomatic AD individuals. The main findings of the study are: (i) **Proteomics** revealed that AsymAD individuals maintain enhanced mitochondrial metabolic functions including BCAA metabolism (ECHS1, BCKDHB), the TCA cycle (IDH2, SUCLA2), fatty acid oxidation (CPT2, ACADSB), and regulatory functions such as mitochondrial translation (MRPL37, TOMM40), structural proteins formation (PDHB, FIS1), and calcium ion transport (MCU, VDAC2) compared to AD; (ii) **Mitochondrial OXPHOS Complexes analysis** showed that most ETC Complexes I-IV required for OXPHOS and ATP generation were preserved in AsymAD and Controls, but were markedly reduced in AD; (iii) **Machine learning models** identified MRPL47, IDH2, CPT2, DBT, MACROD1, and BCAT2 as amongst the top distinguishing mitochondrial features between AsymAD and AD with cross cohort and brain region statistical analysis revealing expression of these proteins in AsymAD to be consistent with Controls but dysregulated in AD; (iv) **Single-nucleus RNA-seq** analysis across neurons and glial cell types demonstrated enhanced expression of mitochondrial genes related to bioenergetics in AsymAD compared to AD, with key genes such as *NDUFV1, BDH1, SLC25A37, SLC25A5, SLC25A25*, and *FTH1* enriched across multiple cell types; (v) **Protein– small molecule interaction analysis** position mitochondrial NAD⁺/NADH-dependent redox pathways as core determinants of bioenergetic stability in AsymAD.

Mitochondrial oxidation of glucose, fatty acids, and amino acids provides metabolic flexibility that is essential for maintaining brain homeostasis ^47^. While AsymAD and AD brains exhibit similar levels of amyloid plaques and tau tangles, the absence of cognitive decline and brain atrophy in AsymAD suggests additional protective mechanisms. Our findings point to mitochondrial protein preservation in AsymAD as a potential driver of this cognitive resilience. While glucose has traditionally been viewed as the brain’s primary energy source, emerging evidence challenges this notion. Recent studies show that neurons, particularly at the synapse, preferentially utilize fatty acids, supporting higher metabolic and enzymatic activity ^48^. This underscores the importance of studying mitochondrial adaptability and fuel switching, which may enable the brain to meet changing energy demands and protect against neurodegeneration. In our cross-cohort, multi-region analyses, the TCA cycle and the ETC consistently emerged as the most preserved mitochondrial pathways in AsymAD compared to AD. The TCA cycle is a central metabolic hub, integrating and linking lipid metabolism, BCAA metabolism, fatty acid oxidation, and ETC activity, all of which are essential for maintaining mitochondrial membrane integrity and sustaining ATP production (**Fig. 6**).

**Figure 6:**
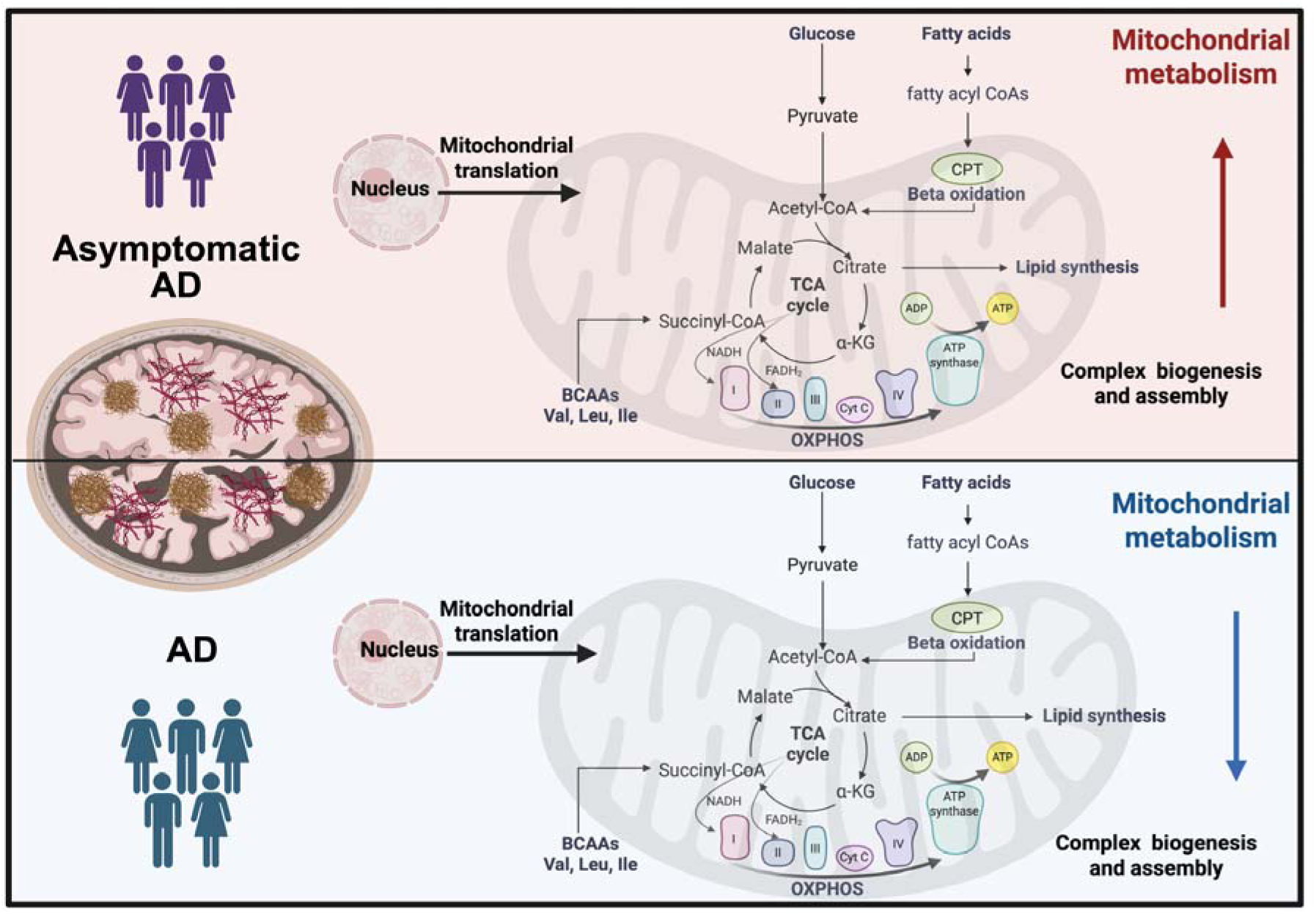
Overview of upregulated mitochondrial regulatory and metabolic pathways in AsymAD compared to AD.

Key proteins consistently enhanced in AsymAD compared to AD include SUCLA2 (Succinate-CoA Ligase ADP-Forming Beta Subunit) and IDH2 (Isocitrate Dehydrogenase 2), both involved in the TCA cycle; CPT2 (Carnitine Palmitoyltransferase 2) and ACADSB (Acyl-CoA Dehydrogenase, Short/Branched Chain), which regulate fatty acid oxidation; and ECHS1 (Enoyl-CoA Hydratase, Short Chain 1) along with BCKDHB (Branched-Chain Ketoacid Dehydrogenase E1 Subunit Beta), both central to BCAA metabolism. Additionally, solute carrier family transporters from the SLC25 family, including SLC25A46 (involved in mitochondrial lipid homeostasis and dynamics) and SLC25A27/UCP4 (essential for neuronal energy homeostasis), were maintained at levels comparable to Controls, supporting their role in sustaining mitochondrial function in AsymAD.

Beyond metabolic enzymes, we also observed preservation of mitochondrial regulatory proteins involved in structural integrity, protein assembly, calcium handling, and translation. These include PDHB (a component of the pyruvate dehydrogenase complex), FIS1 (a fission regulator), and calcium transporters such as MCU (Mitochondrial Calcium Uniporter) and VDAC2 (Voltage-Dependent Anion Channel 2). Notably, members of the mitochondrial ribosomal large subunit (MRPL) family, including MRPL37 and MRPL47, were upregulated in AsymAD. These proteins are essential for mitochondrial translation and critical for maintaining OXPHOS-dependent energy production ^49^. Our study further underscores the importance of investigating mitochondrial biology from a cell type-specific perspective. The majority of mitochondrial functions are governed by nuclear-encoded genes, with over 1,100 mitochondrial genes regulating key aspects of metabolism^50^. Leveraging single-nucleus RNA sequencing (snRNA-seq), we captured the expression of these nuclear-encoded mitochondrial genes across diverse brain cell types, allowing us to dissect their roles in maintaining ETC activity, OXPHOS, the TCA cycle, lipid metabolism, fatty acid oxidation, and the BCAA metabolism in AsymAD vs AD.

We observed a consistent trend of upregulated mitochondrial energy pathways across most cell types in AsymAD, apart from oligodendrocytes and activated astrocytes (Ast CHI3L1). Among all cell types, the ETC function, mitochondrial translation, and lipid metabolism emerged as the most robustly preserved pathways. Some key genes such as *NDUFV1* (*NADH:Ubiquinone Oxidoreductase Core Subunit V1*), *BDH1* (*3-Hydroxybutyrate Dehydrogenase, Type 1*), *FTH1* (*Ferritin Heavy Chain 1*) and solute carrier family SLC25 genes such as *SLC25A37* (iron homeostasis), *SLC25A5* (involved in OXPHOS by coupling ATP production), *SLC25A25* (regulates ATP levels) were enriched across neuronal and glial cell types.

In terms of cell type abundance, AsymAD maintained vulnerable neuronal populations, such as inhibitory PVALB and SST neurons, which are typically reduced in AD^39^. In contrast, microglia and oligodendrocytes were more abundant in AD. The increased mitochondrial activity and expansion of the oligodendrocyte population in AD may reflect a compensatory response to heightened myelination demand in a disease environment characterized by hypomyelination.

Finally, we utilized AD Atlas and DrugBank to identify the pathological associations and small molecules linked to the proteins differentially expressed between AsymAD and AD. NADH emerged as the most highly connected small molecule across 16 candidate protein targets. Early clinical trials using NADH in small human cohorts have showed mixed cognitive outcomes^51–53^ which could be attributed to its poor stability and inability to cross the blood-brain barrier^54,55^. Current strategies now have directed to focus on NAD⁺ (the oxidized form of NADH) precursors such as NR (nicotinamide riboside) and NMN (nicotinamide mononucleotide), which have shown to elevate NAD⁺ levels, restore mitochondrial function, and attenuate Aβ/tau pathology in multiple AD mouse and *C. elegans* models^56^. An ongoing clinical trial is now determining the optimal NR dose for human AD patients (<u>ClinicalTrials.gov ID: NCT05617508</u>). Given that NAD⁺ balance regulates key metabolic and neuroprotective pathways^57^, our finding that NADH-linked targets are central in AsymAD individuals suggests that early therapeutic intervention to sustain mitochondrial bioenergetics via NAD⁺ metabolism may help prevent progression to symptomatic AD, offering a mechanistically distinct alternative to anti-amyloid therapies, which are often associated with adverse effects 8-12. These results highlight the need for more advancing NAD⁺-boosting therapeutics to move toward clinical translation.

Finally, mitochondria are the only organelle in a cell that has its own genome so it can code unique small peptides known as mitochondria-derived micro peptides, including SHLP2, MOTS-c, and SHMOOSE, that have been implicated in protection against neurodegenerative diseases (AD/PD)^58–61^. Among centenarians living beyond 100 years, those carrying APOE4 and who also possessed the mitochondrial-derived peptide humanin P3S performed better on cognitive tests than those lacking this variant. In mouse models of AD with humanized APOE4, treatment with P3S led to a marked reduction in amyloid-β deposition and an increase in OXPHOS genes^62^.

Together, these studies, along with our own findings, underscore mitochondria as a critical organelle capable of influencing whether individuals with AD pathology remain symptom-free, as in asymptomatic AD or progress into the full disease trajectory.

## Data availability

Proteomics and snRNA sequencing used in the study can be obtained from the AD Knowledge portal from the Synapse database(https://www.synapse.org/Home:x). For proteomics data: 4 brain regions were utilized from proteomics, Dorsolateral pre frontal cortex (DLPFC; BA9) data and metadata was obtained from the consensus from 2 cohorts: ROSMAP (Religious Orders Study (ROS) involving older Catholic nuns, priests, and brothers across the United States; Memory and Aging Project (MAP) involving older people across Chicago metropolitan area) and Banner (Banner Sun Health Research Institute, Arizona) (syn25006659 and syn25006658), Para gyrus hippocampus(PHG; BA36) was obtained from Mt Sinai (Mount Sinai School of Medicine Brain Bank, New York; syn25006652 and syn25006648) and pre-motor cortex (BA6) and temporal cortex (BA37) was obtained from ROSMAP (syn25006866 and syn25006867; syn25006835 respectively). For snRNA seq analysis, we leveraged postmortem brain data from ROSMAP cohort for all 6 cell types excitatory, inhibitory, astrocytes, microglia, OPC and oligodendrocytes (syn52293433 and syn3157322).

## Code availability

Codes used for analysis are available in the following GitHub repository: https://github.com/BaloniLab/Asymptomatic_vs_symptomatic_Alzheimers_disease.

## Acknowledgement

This study was supported by grants from the National Institute on Aging (NIA). E.T and P.B were supported by 4RF1AG055549-03. This work was further enabled in part by resources and data provided through the Alzheimer’s Disease Metabolomics Consortium (ADMC) and the Alzheimer’s Gut Microbiome Project (AGMP), funded by the National Institute on Aging under grants U19AG063744, U01AG061359, and R01AG081322 awarded to Dr. Rima Kaddurah-Daouk at Duke University in collaboration with multiple academic institutions. M.A. was supported by the National Institute on Aging through grants U01AG061359, U19AG063744, R01AG069901, U19AG074879, U01AG088562, and R01AG081322. Investigators within the ADMC and AGMP who are not listed as co-authors contributed analysis-ready data but were not involved in study design, data analysis, or manuscript preparation. A full list of ADMC investigators is available at https://sites.duke.edu/adnimetab/team/, and AGMP investigators at https://alzheimergut.org/meet-the-team/. We also acknowledge support from the Mitochondrial Interest Group and the Alzheimer’s Disease Metabolomics Consortium led by R.K.D. for collaborative discussions and data resources.

## Competing Interest Statement

Dr. Kaddurah-Daouk is an inventor on a series of patents on use of metabolomics for the diagnosis and treatment of CNS diseases and holds equity in Metabolon Inc., Chymia, and Metabosensor. M.A. is a co-inventor (through Duke University/Helmholtz Zentrum München) on patents on applications of metabolomics in diseases of the central nervous system; M.A. holds equity in Chymia LLC and IP in PsyProtix and Atai that are exploring the potential for therapeutic applications targeting mitochondrial metabolism in depression. The remaining authors have declared no competing interest.

## Supplementary files

### Supplementary figure legends

**Supplementary Fig 1:** Differentially expressed protein (DEP) abundance calculated after accounting for cell type proportion. a) Number of marker proteins for each cell type. b) Box plot showing the average marker protein expression across individuals (control, AD and AsymAD per cell type). ci-iii) Bar plot showing bias in DEPs obtained between comparisons (AsymAD vs AD; AD vs Control and AsymAD vs Control) before and after cell type correction using Fisher’s exact test per cell type. d & e) Pie chart showing % genes per term for upregulated and downregulated enriched significant pathways across Kegg, Reactome and WikiPathways for AsymAD vs AD respectively using Cytoscape; the Venn diagram shows % DEPs mapped to MitoCarta 3.0. f) Total number of mitochondrial DEPs obtained from 3 methods (Anova, Limma and Limma with correction) across 3 comparisons (AsymAD vs AD, AD vs Control and AsymAD vs Control). g) Venn diagram showing the overlapping genes from 3 analysis for AsymAD vs AD. h) Venn diagram showing the overlapping genes from 3 analysis for AD vs Control.

**Supplementary Fig 2:** Imputation of protein expression matrix and features selected by RFE.

a) Table showing the top 10 proteins with mean and standard deviation shifts after KNN-based imputation. b) Density–histogram plots comparing protein expression distributions before and after imputation for proteins with >15% missing values. c) Schematic summarizing the number of features selected by each classifier and the intersection of common features across classifiers. d) Line plot of features selected by RFECV, showing mean AUC values after 10-fold cross-validation as a function of the number of features. e) SHAP plot displaying the top 20 proteins ranked by absolute mean SHAP values, annotated with their associated pathways. f) Violin plot of SLC25 family proteins identified by RFE across 4 brain regions using *t*-test. g) Violin plot of regulatory proteins identified by RFE, DAP3 and FIS1 across 4 brain regions using *t*-test (ns = *p* > 0.05; * = *p* < 0.05; ** = *p* < 0.01; *** = *p* < 0.001; **** = *p* < 0.0001)

**Supplementary Fig 3:** snRNA seq used to identify cell type mitochondrial gene changes. a) Scatter plot of CERAD score vs Braak score for control, AD and AsymAD individuals, color coded by MMSE score per individual. b) Percentage plot of 14 excitatory cell types clubbed to 3 types and 23 inhibitory cell types to 4. ci-x) Box plots of proportion of each cell type per diagnosis to identify a trend when going from Control AsymAD AD using Jonckheere’s trend test. d) Bar plot of the number of DEGs associated to each pathway, both in up and down direction for 3 comparisons: AD vs Control, AsymAD vs Control and AsymAD vs AD for oligodendrocytes.

**Supplementary Fig 4: Protein–pathway associations of 16 small molecules identified from DrugBank. a)** Bar plot of the number of protein interactors for each of the 16 chemical compounds, derived from DrugBank and BioGRID data. Selected protein names associated with key Reactome pathways are annotated alongside each bar. **b)** Bubble plot illustrate the top five Reactome pathways enriched for each compound, ranked by adjusted p-value. Bubble size reflects the number of associated proteins, while color encodes pathway significance.

**Supplementary tables**: Excel spreadsheet containing tables A-E

## Notes

https://www.synapse.org/Home:x

https://github.com/BaloniLab/Asymptomatic_vs_symptomatic_Alzheimers_disease

