## Supplementary figure for "Mitochondrial bioenergetic signatures differentiate asymptomatic from symptomatic Alzheimer’s disease"

**Supplementary figures**

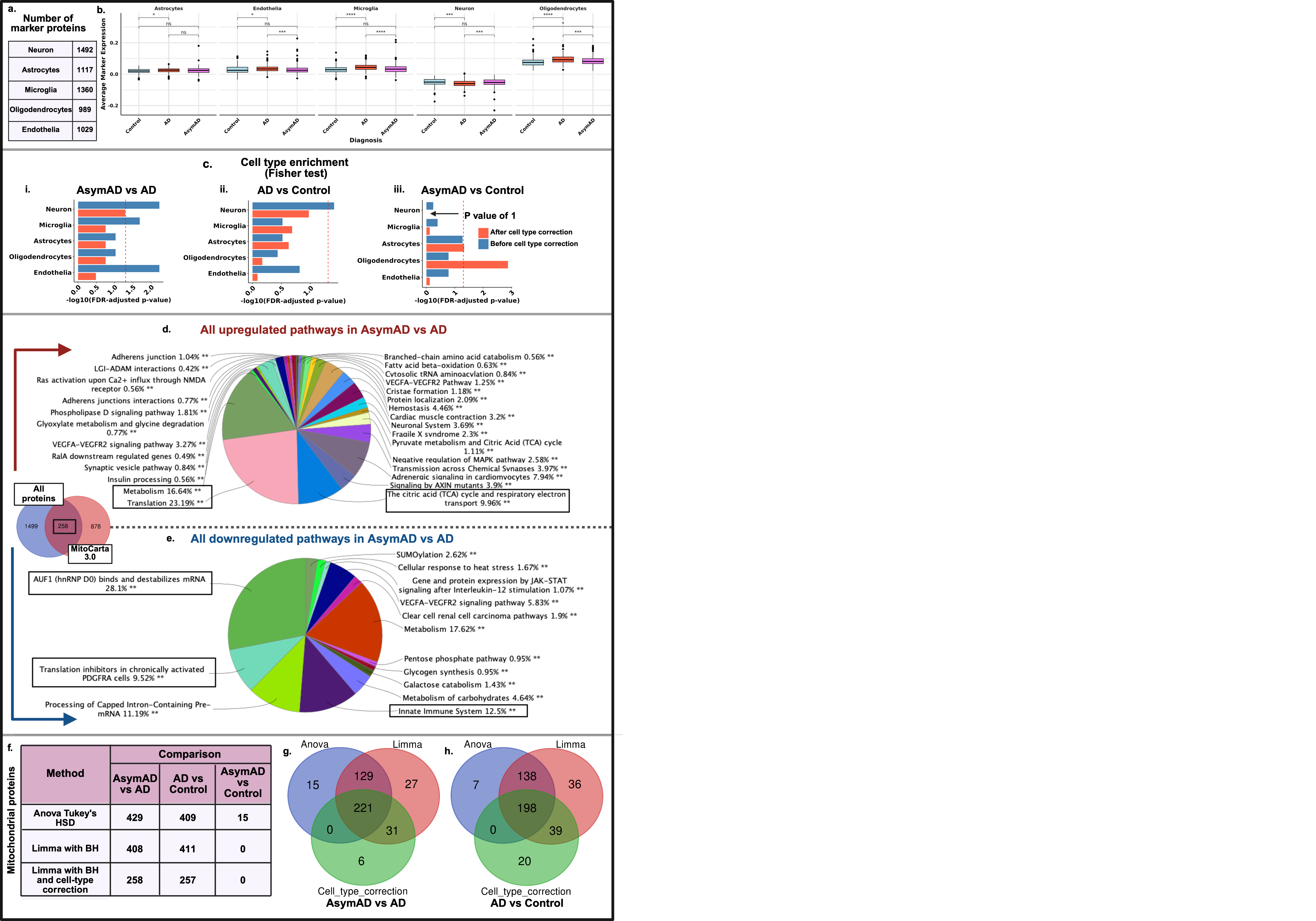

**Supplementary Fig 1:** Differentially expressed protein (DEP) abundance calculated after accounting for cell type proportion. a) Number of marker proteins for each cell type. b) Box plot showing the average marker protein expression across individuals (control, AD and AsymAD per cell type). ci-iii) Bar plot showing bias in DEPs obtained between comparisons (AsymAD vs AD; AD vs Control and AsymAD vs Control) before and after cell type correction using Fisher’s exact test per cell type. d & e) Pie chart showing % genes per term for upregulated and downregulated enriched significant pathways across Kegg, Reactome and WikiPathways for AsymAD vs AD respectively using Cytoscape; the Venn diagram shows % DEPs mapped to MitoCarta 3.0. f) Total number of mitochondrial DEPs obtained from 3 methods (Anova, Limma and Limma with correction) across 3 comparisons (AsymAD vs AD, AD vs Control and AsymAD vs Control). g) Venn diagram showing the overlapping genes from 3 analysis for AsymAD vs AD. h) Venn diagram showing the overlapping genes from 3 analysis for AD vs Control.

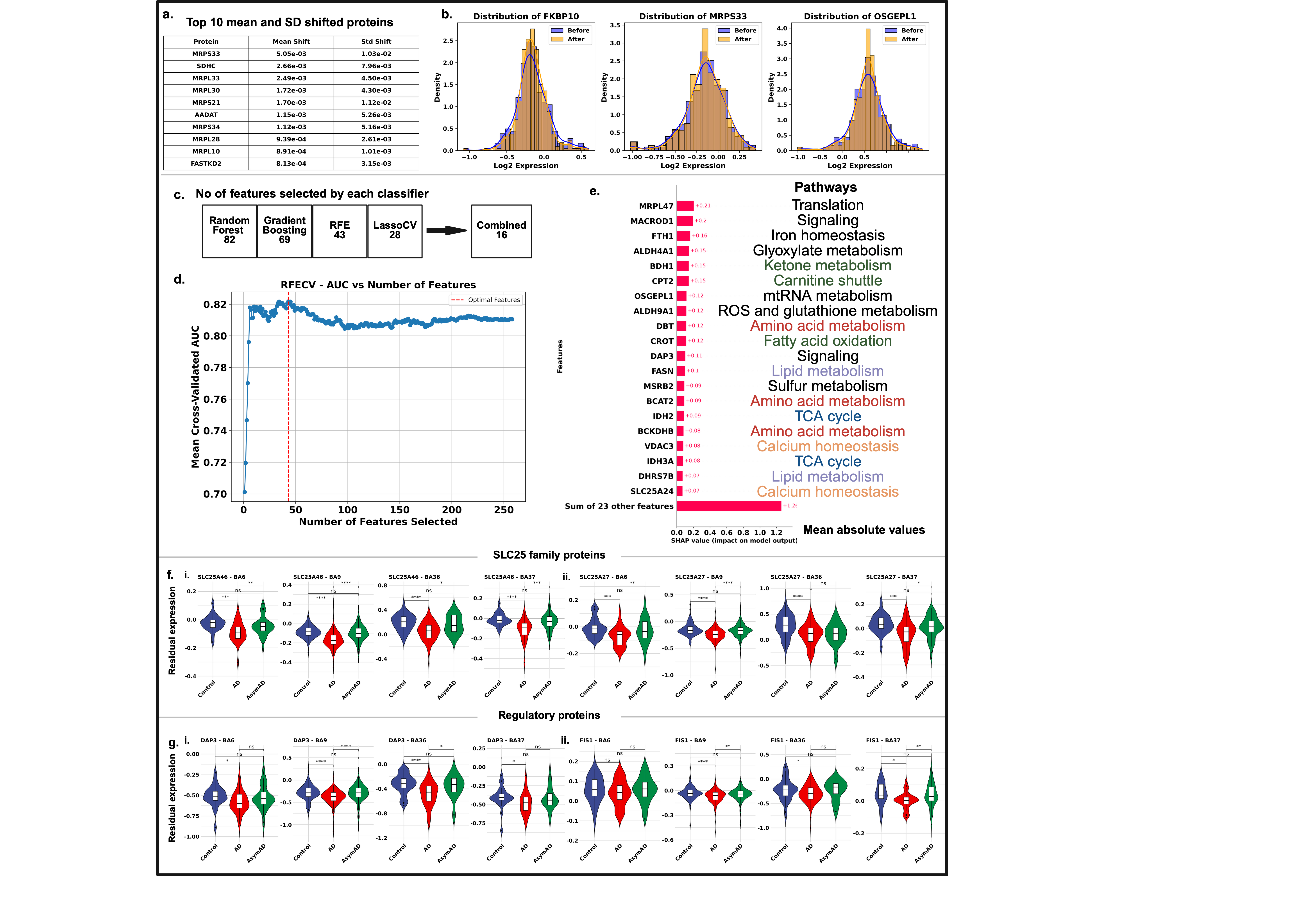

**Supplementary Fig 2:** Imputation of protein expression matrix and features selected by RFE.

a) Table showing the top 10 proteins with mean and standard deviation shifts after KNN-based imputation. b) Density–histogram plots comparing protein expression distributions before and after imputation for proteins with >15% missing values. c) Schematic summarizing the number of features selected by each classifier and the intersection of common features across classifiers. d) Line plot of features selected by RFECV, showing mean AUC values after 10-fold cross-validation as a function of the number of features. e) SHAP plot displaying the top 20 proteins ranked by absolute mean SHAP values, annotated with their associated pathways.

f) Violin plot of SLC25 family proteins identified by RFE across 4 brain regions using *t*-test. g) Violin plot of regulatory proteins identified by RFE, DAP3 and FIS1 across 4 brain regions using *t*-test (ns = *p* > 0.05; * = *p* < 0.05; ** = *p* < 0.01; *** = *p* < 0.001; **** = *p* < 0.0001)

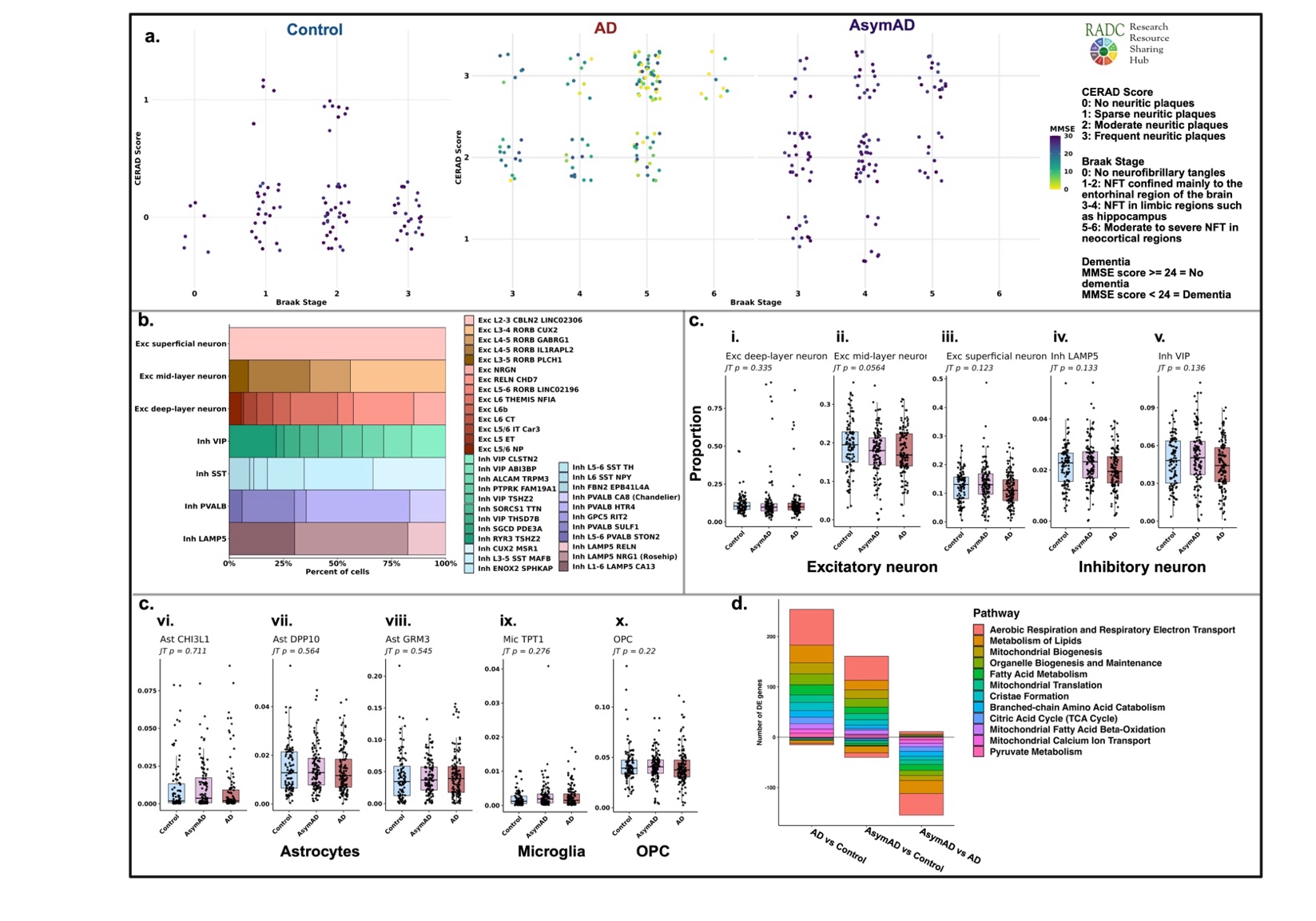

**Supplementary Fig 3:** snRNA seq used to identify cell type mitochondrial gene changes. a) Scatter plot of CERAD score vs Braak score for control, AD and AsymAD individuals, color coded by MMSE score per individual. b) Percentage plot of 14 excitatory cell types clubbed to 3 types and 23 inhibitory cell types to 4. ci-x) Box plots of proportion of each cell type per diagnosis to identify a trend when going from Control
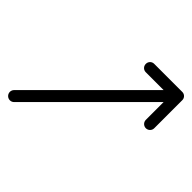
AsymAD
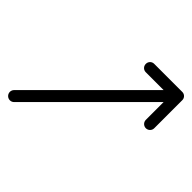
 AD using Jonckheere’s trend test. d) Bar plot of the number of DEGs associated to each pathway, both in up and down direction for 3 comparisons: AD vs Control, AsymAD vs Control and AsymAD vs AD for oligodendrocytes.

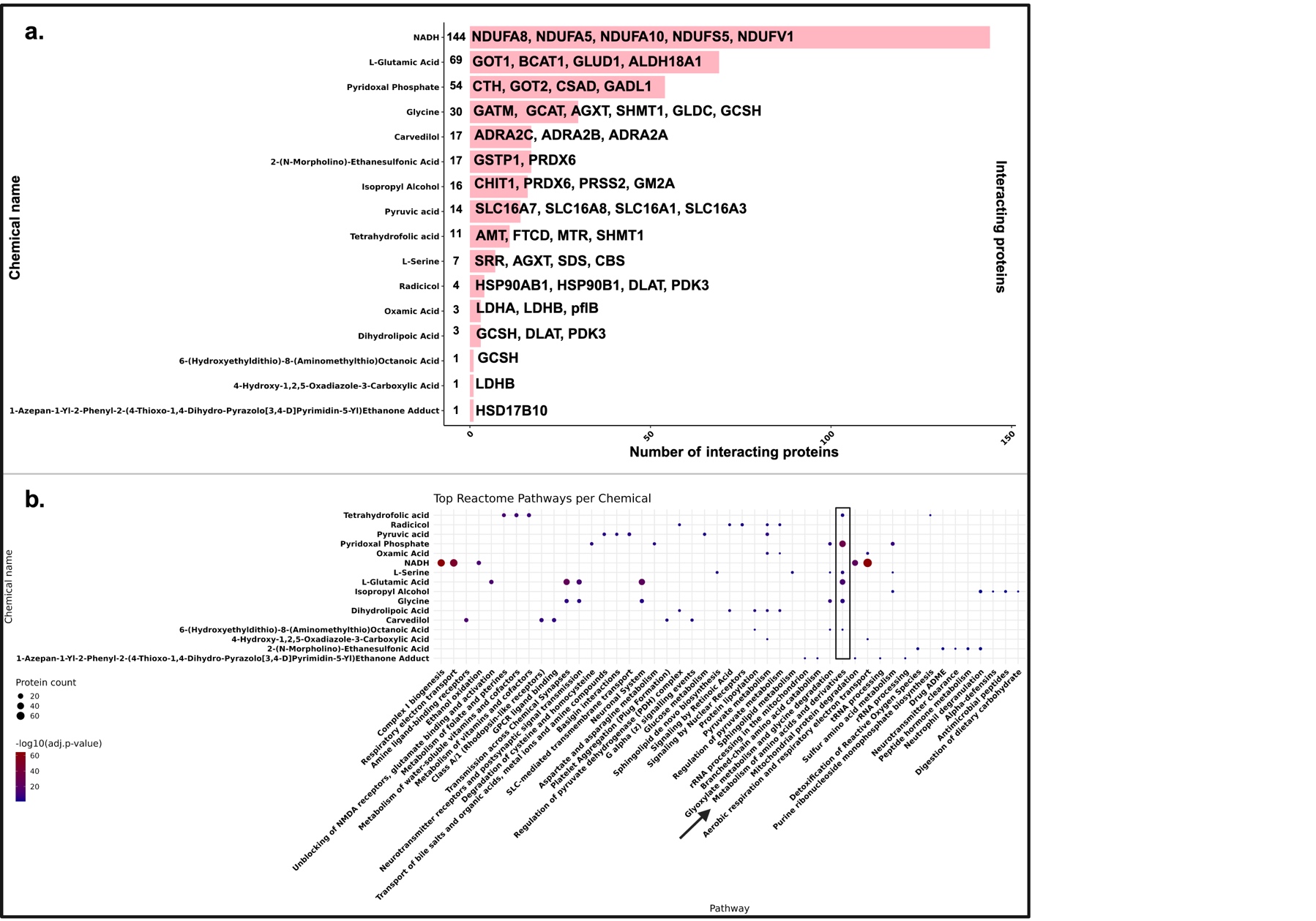

**Supplementary Fig 4: Protein–pathway associations of 16 small molecules identified from DrugBank. a)** Bar plot of the number of protein interactors for each of the 16 chemical compounds, derived from DrugBank and BioGRID data. Selected protein names associated with key Reactome pathways are annotated alongside each bar. **b)** Bubble plot illustrate the top five Reactome pathways enriched for each compound, ranked by adjusted p-value. Bubble size reflects the number of associated proteins, while color encodes pathway significance.
